# Chronic social defeat stress induces a depression-relevant outcome in male prairie voles

**DOI:** 10.1101/2023.10.16.562541

**Authors:** Minerva Rodriguez, Anapaula Themann, Israel Garcia-Carachure, Omar Lira, Alfred J. Robison, Bruce S. Cushing, Sergio D. Iñiguez

## Abstract

**Background:** Stress-induced illnesses, like major depression, are among the leading causes of disability across the world. Consequently, there is a dire need for the validation of translationally-suited animal models incorporating social stress to uncover the etiology of depression. Prairie voles (*Microtus ochrogaster*) are more translationally relevant than many other rodent models as they display monogamous social and parental behaviors and more primate-like neuroanatomy. Therefore, we evaluated whether a novel social defeat stress (**SDS**) model in male prairie voles induces depression-relevant behavioral outcomes.

**Methods:** Adult sexually-naïve male prairie voles experienced SDS bouts from a conspecific pair-bonded male aggressor, 10 min per day for 10 consecutive days. Non-stressed controls (same-sex siblings) were housed in similar conditions but never experienced physical stress. Twenty-four hr later, voles were evaluated in social interaction, sucrose preference, and Morris water maze tests – behavioral endpoints validated to assess social withdrawal, anhedonia-related behavior, and spatial memory performance, respectively.

**Results:** SDS-exposed voles displayed lower sociability and body weight, decreased preference for a sucrose solution, and impairment of spatial memory retrieval. Importantly, no differences in general locomotor activity were observed as a function of SDS exposure.

**Limitations:** This study does not include female voles in the experimental design.

**Conclusions:** We found that repeated SDS exposure, in male prairie voles, results in a depression-relevant phenotype resembling an anhedonia-like outcome (per reductions in sucrose preference) along with social withdrawal and spatial memory impairment – highlighting that the prairie vole is a valuable model with potential to study the neurobiology of social stress-induced depression-related outcomes.

## 1. Introduction

Mood-related illnesses, including major depression, are among the leading causes of disability across the globe (Friedrich, 2017). Despite its high prevalence, the etiology and treatment of depression is not well understood. Depression is difficult to study systematically because it is a complex disease with overlapping and multifaceted syndromes that include anhedonia, altered sleeping and eating patterns, weight fluctuations, memory-related impairment, helplessness, suicidal ideation, and hypothalamic-pituitary-adrenal (**HPA**) axis dysfunction. Underscoring the complexity of this illness, close to 50% of patients suffering from major depression do not respond to most pharmacotherapeutic treatments, accounting for the low remission and recovery rates. Despite this, exposure to social stress, such as bullying, is a well-recognized risk factor for the development of major depression (Swearer and Hymel, 2015). For this reason, identifying the characteristics that precipitate vulnerability using social stress-related animal models (Patel et al., 2019) is critical for uncovering the biological factors that underlie the disease, as well as to establish new approaches for antidepressant medication discovery (Hollon et al., 2015).

Extensive evidence indicates that animals exposed to various forms of stress display ethologically relevant coping behaviors that capture/resemble key features of major depression, including anhedonia, social withdrawal, memory impairment, and HPA axis activation, along with dysregulation of body weight and sleeping patterns (Berton et al., 2012). Although numerous methods have been implemented to stress animals in the laboratory setting, the social defeat stress (**SDS**) approach (Berton et al., 2006), initially introduced in rats and mice as the *resident-intruder* paradigm (Kudryavtseva et al., 1991; Miczek, 1979), is the most validated preclinical model for the study of mood-related illnesses (Hammels et al., 2015). Indeed, when compared to other stress-inducing methods (Atrooz et al., 2021), SDS displays greater face, predictive, and construct validity. For example, SDS-exposed animals not only exhibit reductions in social behavior and preference for sucrose (i.e., anhedonia), but also reflect despair-related behavior when faced with subsequent inescapable stressful situations (i.e., helplessness), memory impairment, and HPA axis overactivation (Hollis and Kabbaj, 2014; Huang et al., 2016; Iñiguez et al., 2014). Furthermore, SDS-induced alterations within the dopaminergic system parallel those observed in postmortem human tissue of individuals who suffered from depression (Der-Avakian et al., 2014; Krishnan et al., 2008; Vialou et al., 2010) with both traditional and fast-acting antidepressant medications reversing behavioral and neurobiological SDS-induced maladaptations (Berton et al., 2006; Donahue et al., 2014; Garcia-Carachure et al., 2020).

While there has been a great interest in the modification and refinement of the SDS model to evaluate stress-induced illnesses (Warren et al., 2020), this experimental approach has been adapted primarily in rats and mice. Yet recently, the monogamous prairie vole (*Microtus ochrogaster*) has emerged as an advantageous model to investigate the biological consequences of stress due to potential higher translational behavioral and neuroanatomical features to humans (McGraw and Young, 2010). For example, unlike most rodent species, prairie voles display monogamous social- and parental-related behavior, with stress altering neurobiological factors and behavior associated with outcomes that resemble anxiety and despair (Lieberwirth et al., 2012; Watanasriyakul et al., 2022). Accordingly, the goal of this investigation is to evaluate whether exposing male prairie voles to the traditional SDS approach results in behavioral and physiological sequalae that are relevant to major depression. We hypothesized that exposing adult male prairie voles to 10-days of SDS, as is commonly done in mice (Berton et al., 2006; Iñiguez et al., 2016), will result in decreases in social behavior, reduced sucrose preference, and spatial memory impairment.

## 2. Materials and Methods

### 2.1 Animals

We used laboratory-reared adult (80-120 days of age) male prairie voles (*Microtus ochrogaster*) that originated from a wild stock trapped near Urbana, IL. Voles were weaned at 21 days of age and housed in same-sex sibling pairs in a colony room maintained on a 14:10 light/dark cycle (light on at 07:00 a.m.) under temperature (∼20-21 °C) controlled conditions and housed in standard polysulfonate rat cages (pine chip bedding) with unrestricted access to food (high fiber rabbit chow) and water (Perry et al., 2019). Experimental procedures were conducted in compliance with the National Institutes of Health *Guide for the Care and Use of Laboratory Animals* (National Research Council (US) Committee for the Update of the Guide for the Care and Use of Laboratory Animals, 2011) and with approval of the *Institutional Animal Care and Use Committee* at The University of Texas at El Paso. Behavioral procedures were performed during the light cycle between 0900-1700 hr.

### 2.2 Experimental Approach

Three separate experiments were conducted to assess whether SDS, a form of social aggression analogous to bullying, would result in depression-relevant behavior in male prairie voles. In the first experiment, male voles were evaluated in the social interaction test 24 hr post stress exposure. In the second experiment, a different cohort of animals were exposed to SDS, and both sucrose preference and social interaction were evaluated 24 hr later. In the last experiment, a different set of animals were tested in the social interaction test 24 hr after defeat, with Morris water maze testing over the next 8 days. Different sets of animals were used across experiments to avoid carryover effects (see Supplemental **Table S1** for experimental groups). Because all animals were screened for sociability after SDS, social interaction data were collapsed across all groups.

### 2.3 Screening for aggressor male voles

Due to their territorial and aggressive behavior toward conspecifics (Carter et al., 1995) pair-bonded and actively breeding male voles were selected as aggressors for the SDS procedure. Specifically, they underwent three consecutive days of aggressive behavior screening to ensure reliable attack latencies before starting the experiment. To do this, the territorialized home-cage (10½ X 19 X 6**⅛**; N40 cage; Ancare, Bellmore, NY) of a pair-bonded couple (male and female) was divided in two separate compartments via a perforated Plexiglass divider. Before aggressive behavior screening, the paired-couple would be physically separated (one on each side of the cage), and the paired-male vole was allowed to acclimate to the opposite side of the cage for 1-min. Following acclimation, a sexually-naïve male screener-vole was placed in the compartment of the paired-male vole, and the time to physically attack the screener was recorded (up to 3 min). Following the 3-min screening session, the screener-vole was removed and checked for potential injuries and returned to its home cage. No behaviors were recorded from the screener vole. Immediately after the screening session was over, the Plexiglass divider was removed from the home cage and the pair-bonded couple was reunified. Male paired-voles with consistent levels of aggressive behavior (attacking within 30 sec) across the 3-day screening were selected as aggressors for the SDS experiments (described below). Close to 50% of paired-voles displayed consistent aggression and were selected as aggressors.

### 2.4 Social Defeat Stress (SDS)

Plexiglass perforated dividers were added to the home cage of pair-bonded and actively breeding aggressors one day prior to SDS, partitioning the home-cage into two equal size compartments (for equipment details see Golden et al., 2011; Iñiguez et al., 2018). For each defeat episode, the pair-bonded male aggressor was moved to the opposite side (left side) of the compartment from their respective pair-bonded female. The pair-bonded female remained in the same compartment throughout the SDS episode (right side), and never physically interacted with the experimental intruder. Sexually-naïve experimental male voles (i.e., intruders) were randomly assigned to experience SDS episodes or were housed in non-stressed control (**CON**) conditions (80-120 days of age). Experimental voles assigned to the SDS condition were placed into the left compartment of the pair-bonded aggressor’s home-cage and exposed to 10-min of physical aggression (**Fig.1A**). Following the 10-min physical confrontation episode, the experimental intruder vole was placed into a holding container for an additional 45-min threat-session with the aggressor (e.g., non-physical sensory interaction; **Fig. 1B**). After the non-physical threat-session, the pair-bonded male aggressor was returned to the right compartment of its home cage to be reunited with its pair-bonded female. The SDS intruder vole was released from the holding container, examined for potential injuries, and remained in the left compartment of the cage for 24 hr until the next SDS episode with a different aggressor. This procedure ensured that all SDS intruder voles were physically defeated by a novel pair-bonded aggressor each day, across 10 consecutive days. CON sexually-naïve male prairie voles were housed in the opposite compartment adjacent to a novel pair-bonded male-and-female couple for 10 consecutive days (but never experienced a physical confrontation with the male pair-bonded aggressor; **Fig. 1C**). Immediately after the last stress episode, all experimental animals (CON and SDS) were single housed in standard laboratory cages (7½ X 11½ X 5; N10 cage; Ancare, Bellmore, NY). Twenty-four hr later, separate groups of experimental voles were tested in a battery of behavioral tasks designed to evaluate social behavior, sucrose preference (anhedonia), or spatial memory performance (described below).

**Figure 1.**
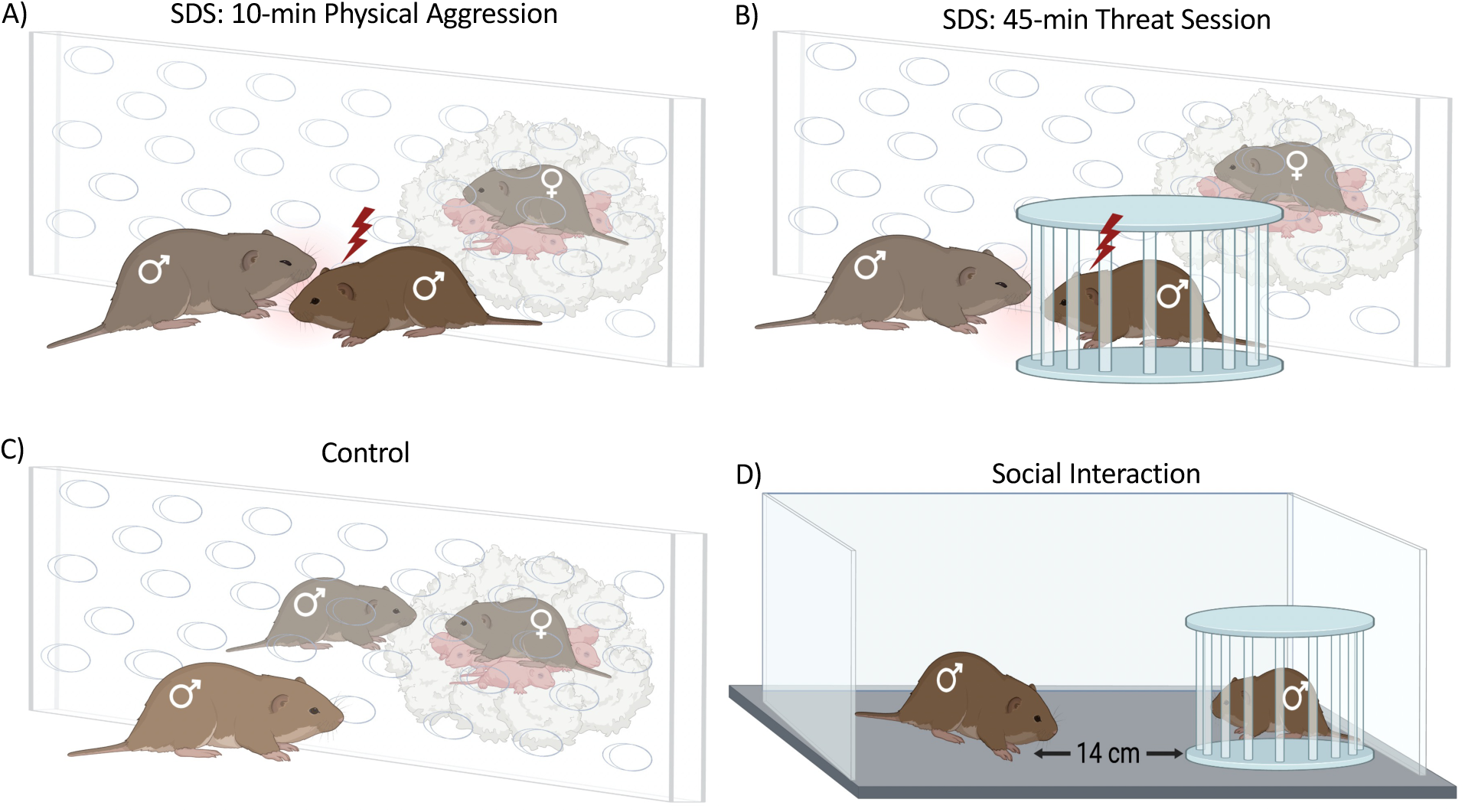
Schematic of the social defeat stress (**SDS**) protocol and social interaction test. (**A**) An experimental vole is placed into the left compartment of a pair-bonded aggressor’s home-cage and is consequently subjected to a physically aggressive encounter for 10 min. (**B**) After the aggression encounter, the SDS-exposed vole is placed within a holding cage for an additional 45-min threat-session with the aggressor. (**C**) Non-stressed control (**CON**) male voles were housed in similar conditions (e.g., the left compartment of a cage with pair-bonded voles); however, they never experienced direct physical- or threat-related encounters. (**D**) Twenty-four hrs after the 10th SDS episode (or CON housing condition), voles were evaluated in the social interaction test. The social interaction zone was the 14-cm area surrounding the circular wire holding cage that contained a novel male social target (not to scale).

### 2.5 Social Interaction Test

To examine the effects of SDS on sociability, all animals were tested in a 2-step social interaction procedure, as described in mice (Iñiguez et al., 2010; Pagliusi et al., 2022). In the first session (2.5 min), the experimental prairie vole was allowed to freely explore an open-field arena (40 cm height x 40 cm length x 40 cm width) that contained an empty circular wire holding-cage (7 cm diameter; Stoelting #60450). After the 2.5 min, the experimental prairie vole was removed from the testing arena (for 30 sec; back in its home cage) and a novel male prairie vole was placed in the wire holding-cage within the testing arena (social target). In the second session (2.5 min) the experimental prairie vole was reintroduced into the testing arena (now containing a social target). Time spent in the social interaction area (defined as a 14 cm circle surrounding the holding cage) was quantified across both sessions (target absent and target present sessions). A social interaction ratio was calculated – where the time (sec) spent in the interaction zone in the presence of the social target was divided by the time spent in the interaction zone in the absence of the social target (for open field schematic see **Fig. 1D** and Iñiguez et al., 2018). As a positive control, distance traveled was recorded during the first 2.5 min (target-absent session) to assess whether locomotor activity or baseline exploratory behavior was influenced by SDS exposure.

### 2.6 Sucrose Preference

To evaluate anhedonia-related behavior, a 2-bottle choice paradigm was adopted (Flores-Ramirez et al., 2018). Here, experimental animals were presented with a choice between consuming water or a sucrose solution. Prairie voles were trained to drink water from two separate bottles across SDS exposures. After the 8^th^ day of SDS, one of the bottles was replaced with a 0.5% sucrose solution. Placement of the bottles (left vs. right) was counterbalanced across cages to avoid potential side preference. A single measurement of water and sucrose consumption was recorded the morning after the last day of defeat stress (0800 hr). Preference for sucrose over water (sucrose/[sucrose + water]) was used as an indicator of reward sensitivity (Iñiguez et al., 2014). Specifically, decreases in preference for sucrose were interpreted as an anhedonia-related phenotype (Willner et al., 1987).

### 2.7 Morris Water Maze (MWM)

The maze was a circular pool with a diameter of 97 cm and a height of 58 cm. The water was filled to a depth of 18 cm, and a standard lamp was used to maintain constant temperature (24±1°C). White nontoxic paint (Handy Art Little Masters Washable Tempera Paint) was added to make the water opaque (Flores-Ramirez et al., 2019; Iñiguez et al., 2012). Spatial cues were positioned on the walls of the testing room, which remained unchanged throughout testing. An escape platform (10 × 10 cm^2^) was submerged to a depth of 0.5 cm in the northeast quadrant throughout spatial acquisition and testing days (but absent during Probe Trial). The MWM procedure consisted of seven phases (Habituation, Spatial training, Test Day, Probe Trial, 1-day break, reversal spatial training, and reversal test day) across 8 days (as depicted in **Figure 4A**). Latency (sec) to reach the platform, swim velocity (cm/sec), and number of platform crossings were recorded via an automated computer tracking system (EthovisionXT, Noldus, Leesburg, VA).

#### 2.7.1 Habituation

Voles were acclimated to the water immersion process (Day 1). Specifically, voles received four shaping trials. On Trial 1, voles were placed on top of a visible escape platform (not immersed) for 10 sec. On Trials 2-4, voles were placed progressively further from the visible escape platform and allowed to swim for up to 60 sec to locate it. If the vole failed to find and mount the platform within the time allotted, the experimenter would gently guide the vole to the platform. No data was collected during the habituation phase.

#### 2.7.2 Spatial Training

Spatial training consisted of two days (Days 2-3), with the escape platform located in the northeast quadrant of the water maze. Prairie voles underwent nine spatial training trials per day, totaling 18 independent spatial training trials. On each trial, the prairie vole was released from one of three starting/release points, located between the northwest, southwest, and southeast quadrants. Voles were given 60 sec to locate the submerged escape platform per training trial. Voles were released three times from each of the three different starting points in a randomized fashion ensuring that they did not start from the same release point on two consecutive trials. If the vole failed to find the escape platform within 60 sec, the experimenter gently guided the animal to the platform, where it was allowed to remain for 10 sec. After each trial, voles were dried with a towel and placed on a temporary holding cage for a 45-sec intertrial period under a standard heat lamp.

#### 2.7.3 Test Day

Twenty-four hr after spatial training (Day 4), voles were tested on a single memory retention trial. Here, each vole was released from the same starting point (southwest; farthest from platform) and was allowed up to 60 sec to swim and locate the platform. Longer latency (sec) to find the escape platform was interpreted as impaired memory performance.

#### 2.7.4 Probe Trial

Twenty-four hr after the Test Day (Day 5), voles underwent a single probe trial wherein the escape platform was removed from the pool. Voles were released from the same starting point (southwest, farthest from the quadrant that used to contain the platform) and were allowed to freely swim for 60 sec. Number of crossings within the area that used to contain the platform were recorded – lower number of crossings were interpreted as impaired spatial memory performance.

#### 2.7.5 Reversal Spatial Training and Test Day

To increase the demands of the memory task, a reversal 1-day spatial training protocol was conducted 48 hr following the Probe Trial (Day 7). Here, the escape platform was relocated to the opposite quadrant from its initial position (from northeast to southwest quadrant). Voles underwent nine training trials to learn the new location of the platform within a single day. Specifically, animals were released 3-times from each of the three separate starting-points in a randomized fashion to ensure that they never started from the same point on two consecutive trials. A single memory retention trial (i.e., Reversal Test Day) was conducted 24 hr after the reversal training day (Day 8), wherein the submerged escape platform was located on the southwest quadrant of the maze. Voles were released from the same starting point (northeast) and allowed up to 60 sec to swim and locate the platform. Longer latency to locate the platform was interpreted as memory impairment.

### 2.8 Statistical Analyses

Voles were randomly assigned to the different stress conditions (CON or SDS) and behavioral endpoints (Supplemental Table S1). Data were analyzed adopting mixed analyses of variance (**ANOVA**) techniques, with stress (between variable), swim trial (repeated measure), and day of weight recording (repeated measure) as sources of variance. Post hoc comparisons were analyzed using Dunnett’s tests. When appropriate, independent two-group comparisons were evaluated with a Student’s t-test. Data are presented as mean ± standard error of the mean (**SEM**) and statistical significance was defined as p≤ 0.05. Analyses were performed using IBM SPSS statistical software (International Business Machines Corporation, Armonk, NY; version 28). See **Supplemental Table S2** for statistical analyses values.

## 3. Results

### 3.1 Social defeat stress decreases social behavior

**Figure 2** displays the effects of SDS on social behavior in adult sexually-naïve male prairie voles (N= 58). When compared to CONs (n= 29), voles exposed to SDS (n= 29) displayed decreases in social interaction ratios (t_56_= 2.21, p< 0.05, Fig. 2A). When assessing locomotor activity (Fig. 2B) during the no-target present trial (2.5 min), no differences were observed between the two experimental conditions (t_56_= 0.29, p> 0.05).

**Figure 2.**
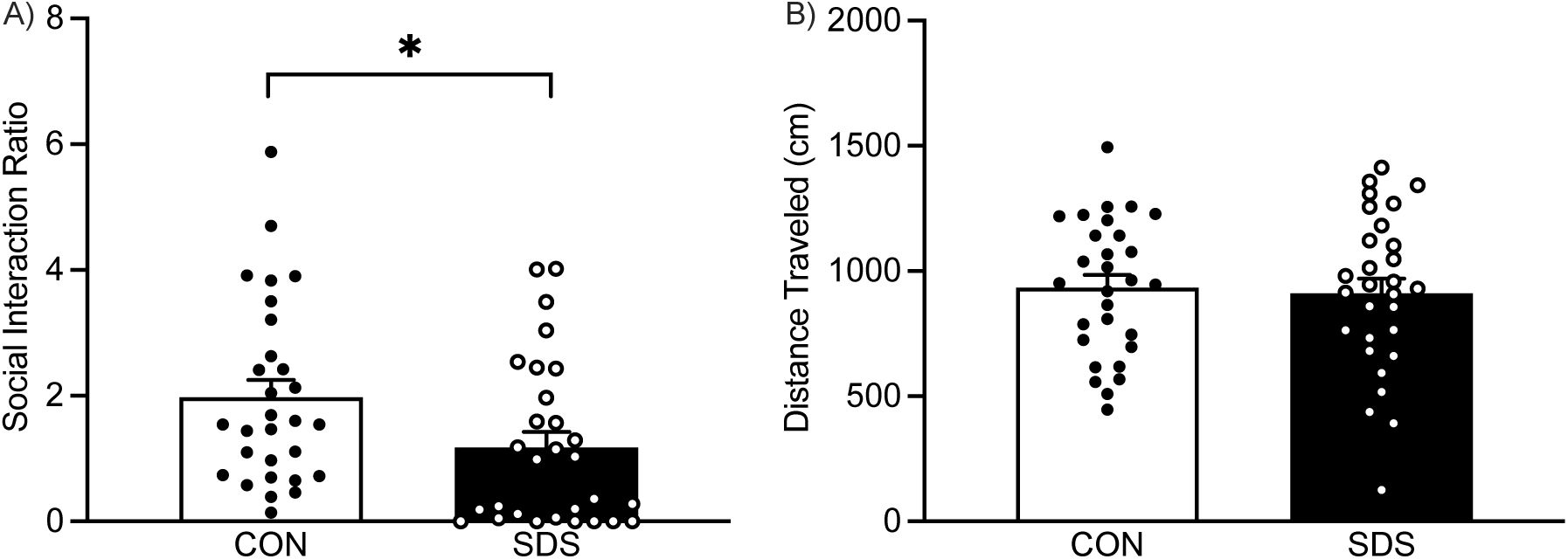
Effects of social defeat stress (**SDS**) on social behavior in adult male voles. (**A**) When compared to the non-stressed control (**CON**) group (n= 29), the SDS voles (n= 29) displayed lower interaction ratios 24 hrs after the 10^th^ defeat episode. (**B**) No differences in locomotor activity (distance traveled in cm) during the first 2.5 min session of the social interaction test (target-absent session) were detected between experimental conditions. Data are presented as mean + SEM. *p< 0.05.

### 3.2 Social defeat stress decreases sucrose preference

**Figure 3** shows the effects of SDS (n= 10) and CON (n= 10) housing conditions on a 2-bottle choice sucrose preference test. SDS-exposed voles displayed a significant decrease in preference for a 0.5% sucrose solution when compared to non-stressed CONs (t_18_= 1.71, p≤ 0.05; Fig. 3A). No differences in total liquid intake (water + sucrose) were noted between the experimental conditions (t_18_= 0.97, p> 0.05; Fig. 3B).

**Figure 3.**
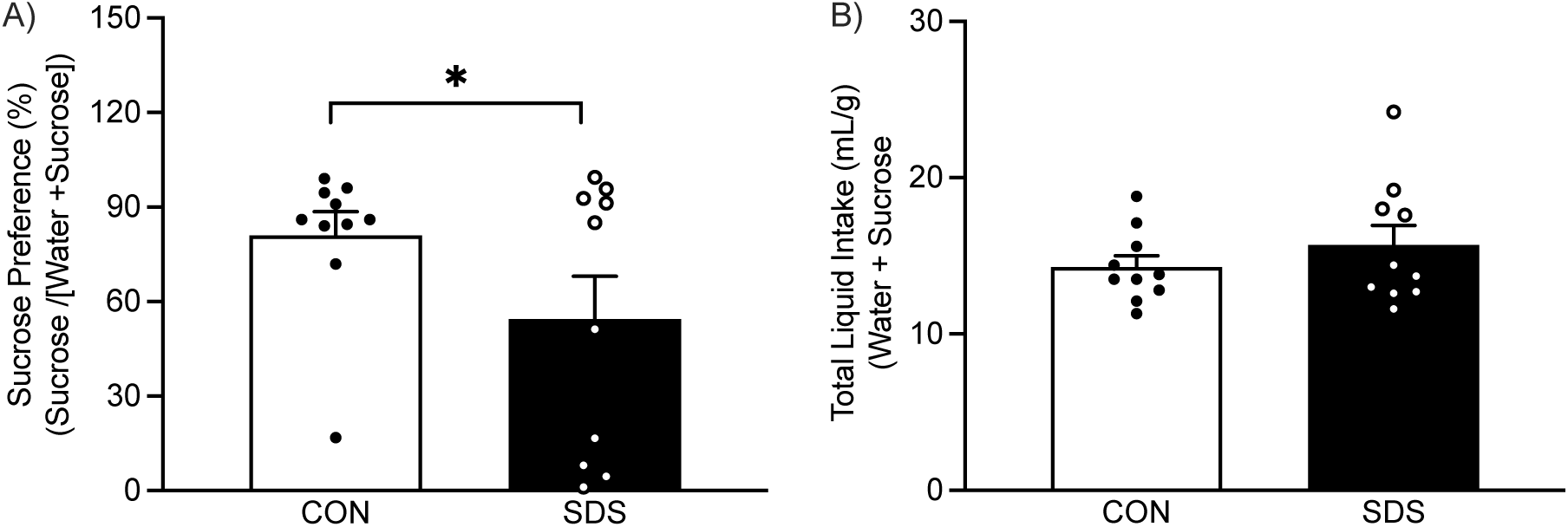
Effects of social defeat stress (SDS) on sucrose preference in adult male voles. (**A**) Twenty-four hrs after the last defeat episode, SDS voles (n= 10) displayed decreased preference for a 0.5% sucrose solution, when compared to the non-stressed control (CON; n= 10) group. (**B**) No differences in total liquid intake (sucrose + water) were observed between the experimental groups. Data are presented as mean + SEM. *p< 0.05.

### 3.3 Social defeat stress impairs spatial memory performance on the MWM

The effects of SDS on a MWM spatial acquisition task are shown in **Figure 4**. Here, a 2-way mixed-design ANOVA with stress (between group variable) and swim trial (within group variable) as sources of variance revealed that the latency (sec) to locate the platform was influenced by a swim trial main effect (F_17, 204_= 1.94, p< 0.05), but not stress exposure (F_1,12_= 1.33, p> 0.05) or their interaction (F_17, 204_= 1.64, p> 0.05). *Post hoc* comparisons indicate that, when compared to Trial 1, voles were taking less time to locate the platform on Trials 8, 9, 16, 17, and 18 (p< 0.05, respectively; **Figure 4B**). One day after spatial training, SDS voles displayed increases in the time to locate the escape platform on Test Day (**Fig. 4C**), however, this difference did not reach statistical significance (t_12_= 1.07, p> 0.05). Yet, during a standard probe trial (one day later), SDS-exposed voles displayed significantly fewer crossings in the area that previously contained the platform, when compared to CONs (t_12_= 2.94, p< 0.05, **Fig. 4D**), uncovering spatial memory deficiencies. Thus, to increase the demands of the memory task, a single reversal spatial acquisition session was conducted 48 hr post the Probe Trial. **Figure 4E** shows that the time to locate the new platform location (southwest) was influenced by a swim trial main effect (F_8,96_= 4.08, p< 0.05), but not stress exposure (F_1,12_= 1.53; p> 0.05) nor their interaction (F_8,96_= 0.95, p> 0.05). Here, voles displayed a decrease in the time to locate the platform on the last five swim trials (Trials 5-9) when compared to Trial 1 (p< 0.05, respectively) – thus, highlighting acquisition of the spatial task across the 9 swim trials. Twenty-four hr later, on the Reversal Test Day (**Fig. 4F**), SDS-exposed voles exhibited longer time to reach the location of the escape platform when compared to CONs (t_12_= 2.50, p< 0.05), once again, uncovering SDS-induced spatial memory-related deficits.

**Figure 4.**
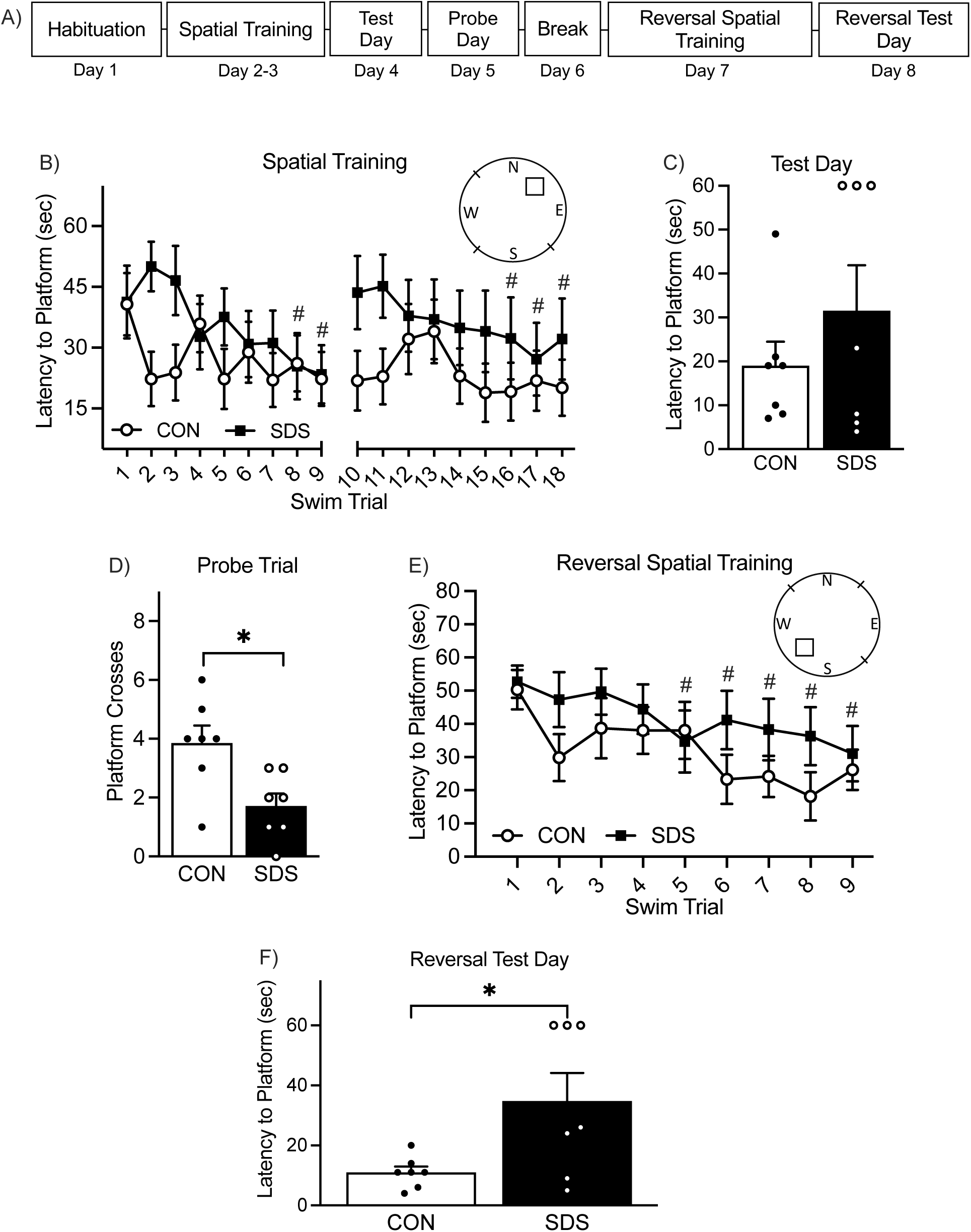
Effects of social defeat stress (**SDS**) on Morris water maze performance in adult male voles. (**A**) Experimental timeline depicting spatial training sessions and memory retention trials between adult male voles exposed to SDS or non-stress control (**CON**) housing conditions. (**B**) During the initial 2-day spatial training acquisition sessions, all voles learned to locate the submerged escape platform (on northeast quadrant) independent of stress exposure. When compared to the 1^st^ swim trial, all voles displayed decreases in latency (sec) to find the platform on trials-8, 9, 16, 17, and 18 (^#^p< 0.05, respectively). (**C**) Twenty-four hr after the spatial training trial (Test Day), no differences in the latency to find the platform were noted between SDS (n= 7) and CON voles (n= 7). (**D**) The following day (Probe Day), SDS voles displayed fewer crossings within the area that previously contained the escape platform when compared to CONs (\**p*< 0.05). (**E**) During a single-day reversal spatial training acquisition session consisting of 9-trials, all voles learned to locate the new location (southwest quadrant) of the escape platform independent of stress exposure. Specifically, when compared to Trial 1, voles displayed decreases in the latency to find the platform on Trials 5-9 (^#^p< 0.05). (**F**) Twenty-four hr after reversal training (Reversal Test Day), SDS voles displayed longer latency (sec) to locate the escape platform when compared to CONs (*p< 0.05). Data are presented as mean + SEM. *p< 0.05 when compared with male CON group. ^#^p< 0.05 when compared to Trial-1. **N**, north; **E**, east; **W**, west; **S**, south. Dashes on circular water maze schematic represent the 3 different starting release points.

### 3.4 Social defeat stress does not influence swimming velocity on the MWM

**Figure 5** displays the effects of SDS on swim velocity across the different phases of the MWM procedure. During the initial 2-day spatial training, a mixed 2-way ANOVA indicated that swim velocity (cm/sec) was influenced by swim trial (F_17, 204_= 2.26, p< 0.05), but not stress (F_1,12_= 0.27, p> 0.05) or their interaction (F_17, 204_= 0.55, p> 0.05). *Post hoc* tests indicated that, when compared to Trial 1, voles were swimming slower on Trials 9, 15, 16, and 18 (p< 0.05, respectively; **Fig. 5A**). No differences in swim velocity were noted on Test Day (t_12_= 0.03, p> 0.05; **Fig. 5B**) or the Probe Trial (t_12_= 0.52, p> 0.05; **Fig. 5C**) as a function of SDS. During the reversal spatial acquisition session day, no differences in velocity were detected as a function of swim trial (F_8,96_= 0.94, p> 0.05), stress (F_1,12_= 0.64, p> 0.05) or their interaction (F_8,96_= 1.13, p> 0.05, **Fig. 5D**). Lastly, no differences in velocity were noted between the experimental groups (t_12_= 0.88, p> 0.05) during the Reversal Test Day (**Fig. 5E**).

**Figure 5.**
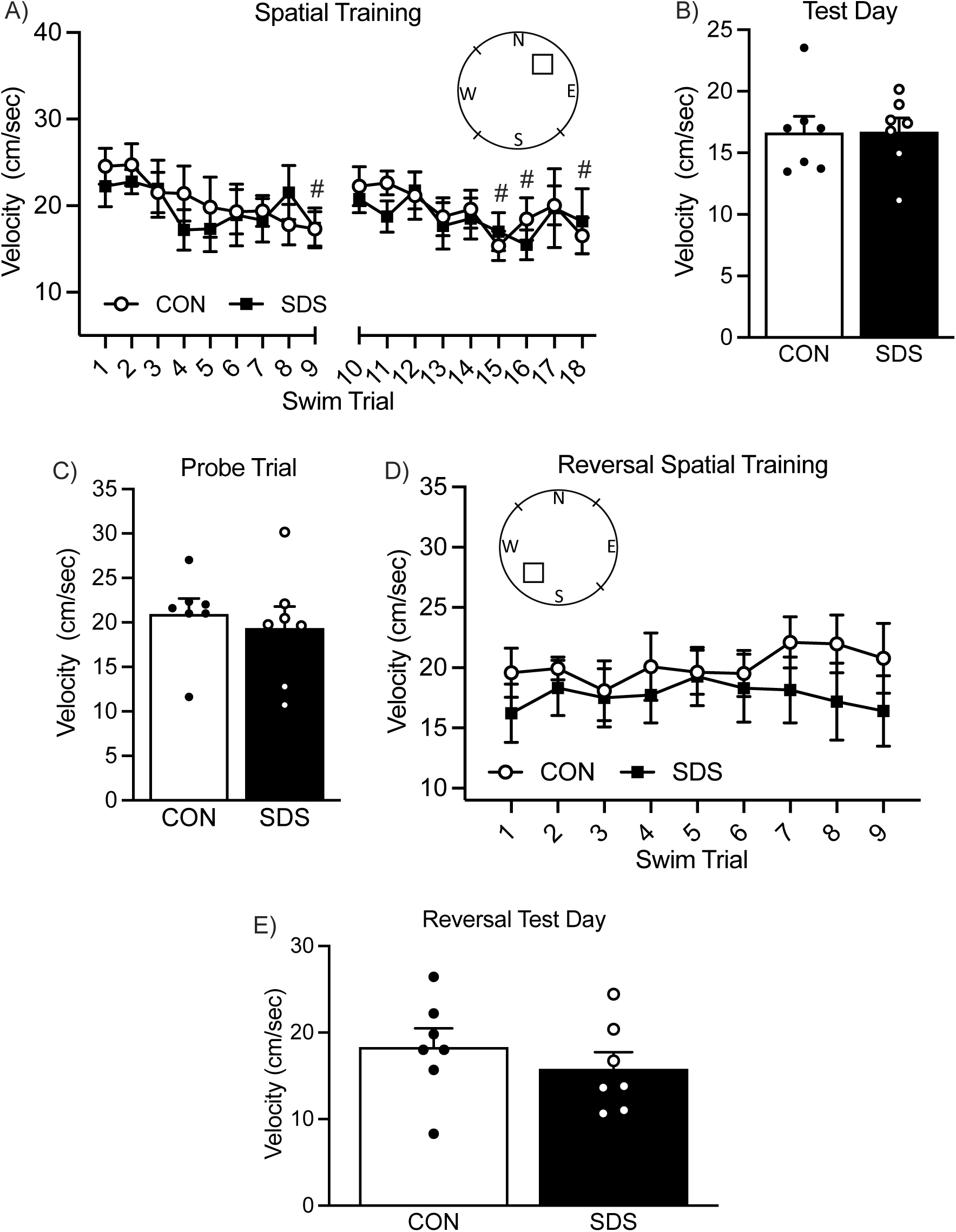
Effects of social defeat stress (**SDS**) on swimming velocity (cm/sec) in the Morris water maze in adult male voles. (**A**) During the initial 2-day spatial training session (Trials 1-18), non-stressed control (**CON**) and SDS voles displayed similar swimming velocity across training trials. However, there was a decrease in velocity as trials progressed. Specifically, when compared to the 1^st^ swim trial, voles displayed a decrease in velocity on Trial 9, as well as in the last 4 swimming trials (Trials 15-19; ^#^p< 0.05, respectively). Likewise, SDS did not influence swim velocity during the (**B**) Test Day or (**C**) Probe Trial. (**D**) When voles returned to the water maze for a single day Reversal Spatial Training session of 9 trials, no differences in swimming velocity were observed between the groups. (**E**) Lastly, no differences in swim velocity were noted as a function of SDS on the reversal test day. Data are presented as mean + SEM. ^#^p< 0.05 when compared to Trial 1. **N**, north; **E**, east; **W**, west; **S**, south. Dashes on circular water maze schematic represent the 3 different starting release points.

### 3.5 Social defeat stress decreases body weight

The effects of SDS on body weight change are presented in **Figure 6**. Body weight change was calculated by subtracting the total weight of the experimental animal starting on the initial day of defeat (or CON condition) to each subsequent day; thus, a positive number indicates weight increase, whereas a negative number indicates weight decrease (Warren et al., 2013). Here, a two-way mixed ANOVA with stress (between group variable) and day (repeated measure) as sources of variance indicated that weight change was influenced by stress exposure (F_1,18_= 8.89, p< 0.05), day (F_10,180_= 23.64, p< 0.05) and their interaction (F_10,180_= 3.76, p< 0.05). Specifically, *post hoc* comparisons revealed that SDS-voles displayed a reduction in weight change when compared to CONs from day-3 to day-10 (p< 0.05). This defeat-induced reduction of body weight remained 24 hr after the last SDS episode (t_18_= 3.26, p< 0.05), right before social interaction testing.

**Figure 6.**
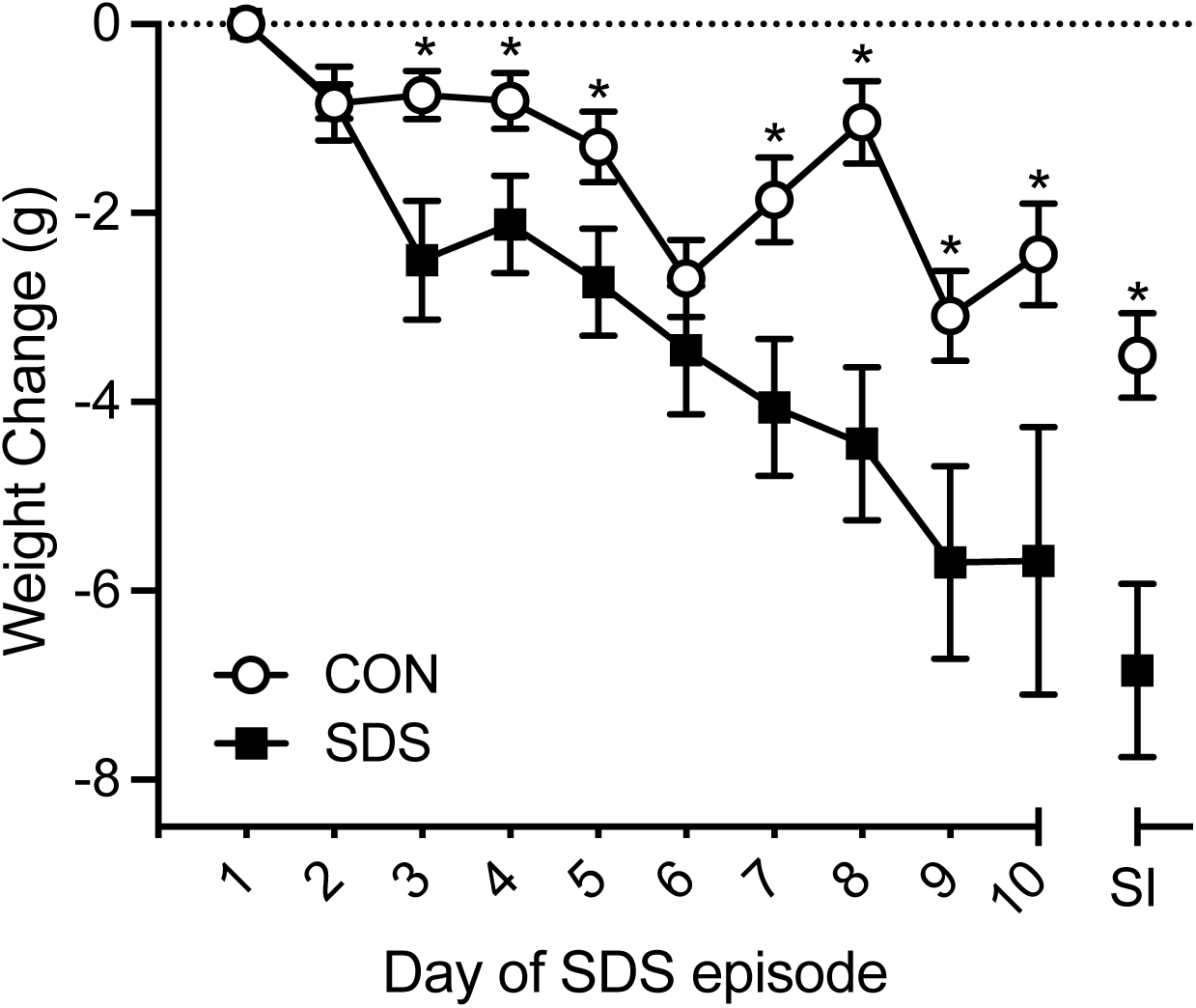
Effects of social defeat stress (**SDS**) on body weight-change in adult male voles. When compared to the non-stressed control (**CON**) group (n= 10), voles exposed to SDS (n= 10) displayed decreases in body weight-change as of day 3 of defeat. This SDS-induced decrease in body weight-change remained 24 hr after the 10^th^ exposure, prior to social interaction (**SI**) testing. Data are presented as weight change in grams + SEM. *p< 0.05.

## 4. Discussion

Preclinical stress models have been instrumental in our understanding of the neurobiology of depression- and antidepressant-relevant behavior (Krishnan and Nestler, 2008; Warren et al., 2020). The SDS model is among the most validated approaches for the study of social stress-induced illnesses, but most work implementing SDS has been conducted across different strains of rats and mice (Burke and Miczek, 2015; Duque-Wilckens et al., 2020; Iñiguez et al., 2014; Newman et al., 2019) which capture some, but not all, human depression-relevant syndromes (Berton et al., 2012). The prairie vole displays behavioral responses to various types of stress similar to those shown by mice and rats (Donovan et al., 2018; Donovan et al., 2020) while exhibiting monogamous and parental-like behavior (Young et al., 2011), thus potentially providing additional translational opportunities for the study of stress-induced illnesses. For this reason, we evaluated whether exposing adult male voles to the SDS paradigm (Berton et al., 2006) would result in behavioral outcomes that recapitulate social withdrawal, anhedonia, and spatial memory impairment – key syndromes exhibited in patients suffering from major depression.

After 10 episodes of SDS, we found that adult male defeated voles displayed reductions in body weight (**Fig. 6**) and social behavior (by spending less time in the interaction zone when compared to CONs; **Fig. 2**). These stress-induced outcomes complement what has been recently reported in adolescent male prairie voles (Sailer et al., 2022) – which display increases in time to approach a social target after a similar 10-day SDS protocol. Likewise, both female prairie and mandarin voles exhibit decreases in sociability after SDS exposure, however, additional depression-related phenotypes associated with helplessness, per increases in immobility in the forced swim and tail suspension tests, are observed only after chronic (14 episodes; Wang et al., 2019) and not acute SDS protocols (3 episodes; Tickerhoof et al., 2020). Together, these studies indicate that repeated SDS episodes (≥10) are needed to capture a robust depression-relevant profile (social withdrawal and learned helplessness) in this monogamous rodent. Chronic versus acute stress exposure is an important factor to consider when validating animal models for the study of mood-related disorders, because humans, as well as animals, do not often develop depressive-related phenotypes after acute exposure to stress; thus, repeated defeat exposures provide stronger face validity to evaluate/characterize behavioral endpoints designed to recapitulate chronic stress-induced depression-related syndromes (Tran and Gellner, 2023). Indeed, SDS-induced reductions of social behavior, with 10 SDS exposures, are traditionally adopted as a depression-relevant endpoint across the preclinical literature, because both chronic administration of selective serotonin reuptake inhibitors (Krishnan et al., 2008) as well as acute exposure to ketamine (Donahue et al., 2014; Garcia-Carachure et al., 2020; Iñiguez et al., 2018) reverse the enduring social dysfunction reported after chronic episodes of stress. Moreover, given the wide distribution of social interaction ratios after SDS (**Fig. 2**), future work is needed to delineate a formal way to categorize susceptible versus resilience-related behavior within the SDS-vole population, as has been done with mice (Krishnan et al., 2007; Vialou et al., 2010). This may be important because resilience underlies active neurobiological processes that are different from non-stressed animals (Feder et al., 2009; Pagliusi et al., 2022). Also, future pharmacological approaches evaluating whether the administration of traditional and/or fast-acting antidepressant medications rescue the SDS-induced maladaptive social behavior observed in prairie voles are needed. This is necessary to directly provide predictive/pharmacological validity to the vole SDS model for the study of mood-related disorders.

In addition to dysregulation of social behavior (Li et al., 2023), most rodents subjected to SDS display reward-related deficits (Peartree et al., 2012), exhibit alterations in dopaminergic activity across different brain regions (Iñiguez et al., 2016; Jin et al., 2015; Slattery and Cryan, 2017), and show reduced sucrose preference – commonly referred to as an anhedonia-related response (i.e., the decreased ability to experience pleasure). Therefore, we also evaluated if changes in sucrose preference would be evident after 10-days of SDS. Not surprisingly, SDS-exposed voles displayed a lower preference for a sucrose solution, without changes in total liquid intake (**Fig. 3**). This finding is consistent with prior vole studies implementing chronic stress approaches resulting in reductions of sucrose preference after 4-weeks of social isolation (Grippo et al., 2008). Of note, previous work adopting acute stress protocols do not report anhedonia-like responses nor increases in corticosterone (Arai et al., 2016; Tickerhoof et al., 2020) – highlighting that chronic exposure to SDS is necessary to uncover anhedonia-like states, social withdrawal, and helplessness-related behavior in prairie voles (Sailer et al., 2022).

Across the clinical literature, exposure to various forms of stress impairs spatial memory performance (Brown et al., 2020; Lupien et al., 2005) comparable to patients suffering from major depression (Gould et al., 2007). Thus, we also opted to evaluate the effects of SDS on MWM performance; a task validated for the assessment of spatial memory in rodents (Novick et al., 2013), including voles (Tomas Pereira and Burwell, 2015). Here, we found that SDS-voles displayed reduced spatial memory when evaluated on a standard probe trial (48 hr post spatial navigation training; **Fig. 4D**). Furthermore, when increasing the demands on the spatial learning/memory task by subjecting voles to a single-day reversal training session, we found that defeated voles spent longer time to locate the escape platform on the reversal testing day (**Fig. 4F**), thus, revealing spatial memory retrieval deficits. Importantly, no differences in swimming velocity were noted as a function of stress across the MWM procedure (**Fig. 5**), emphasizing that performance was due to stress-induced memory impairment and not changes in general locomotor activity (**Fig. 2B**) or body weight fluctuations (**Fig. 6**). Collectively, we report for the first time that 10-days of SDS results in spatial memory deficits, along with decreases in body weight and social behavior, as well as lowered sucrose preference in male prairie voles, establishing that monogamous prairie voles, like rats and mice, display a robust depression-relevant behavioral profile after 10-days of SDS.

### 4.1 Limitations

A limitation of the present study is that we did not evaluate the effects of SDS on depression-relevant endpoints in female voles, consequently limiting the translatability of the present findings to males only. Women, when compared to men, are more likely to be diagnosed with major depression, and thus it will be necessary to evaluate whether repeated exposure to SDS mediates a depression-relevant outcome as a function of sex. Of note, previous work suggests that a brief episode of social conflict increases corticosterone in female prairie voles (Smith et al., 2013). For this reason, it is likely that 10 bouts of SDS will result in depression-relevant outcomes in female voles, particularly because 3-SDS exposures induce an anxiogenic-related effect (Tickerhoof et al., 2020) and prior work in mice further indicates that higher number of SDS episodes (<10) mediates a robust depression-related phenotype (Bondar et al., 2018). Importantly, because both male and female voles display aggression towards conspecifics of either sex, this preclinical model can be modified/refined to induce social conflict in an all-female and/or mixed setting, theoretically allowing researchers to dissect the deleterious effects of chronic social stress on depression-relevant behaviors as a function of sex and/or age. Accordingly, the SDS vole model may provide a platform to uncover the specific role such risk factors play in mood-related psychopathology and discovery of novel interventions (Bath et al., 2023; Wells et al., 1985).

## 5. Conclusion

Overall, our data indicate that implementing the SDS paradigm in adult male prairie voles, as is commonly done in mice (Chandra et al., 2017; Francis et al., 2015), captures a depression-related behavioral outcome resembling social dysfunction, anhedonia, and spatial memory impairment. This collective depression-relevant profile provides a platform for the utilization of prairie voles to expand our understanding of the underlying mechanisms of chronic stress-induced mood-related phenotypes. This could be particularly important because prairie voles share additional characteristics that resemble human behavior (monogamy and parental-related behavior) when compared to other rodents. As such, this preclinical model has the potential to provide a more translationally-relevant approach to uncover the behavioral and molecular underpinnings of stress-induced mood-related illnesses like major depression.

